# Sex-dependent carry-over between winter associations and breeding proximity in a communally roosting parrot

**DOI:** 10.64898/2025.12.21.695283

**Authors:** Julia Penndorf, Annegret Csak, Lucy M Aplin

**Author notes:** corresponding author: Julia Penndorf.

## Abstract

Between-season carry-over in social associations is widely observed and can have important benefits for individuals, ranging from reduced conflict between neighbours to cooperative nest defence. Yet the underlying drivers of social carry-over effects remain unclear, as current research has predominantly focused on species where breeding territories are influenced by individuals’ non-breeding spatial distribution, making it challenging to disentangle social and spatial preferences. Sulphur-crested cockatoos (*Cacatua galerita*) form communal night-roosts with fission-fusion dynamics around the roost-site. During spring, breeding individuals split from the roost to occupy nearby tree hollows, but remain non-territorial, allowing us to test whether individuals exhibit a preference to breed in proximity to winter social associates. We find that male—but not female— winter social associations are correlated with breeding location; males bred closer to their winter associates, and more socially connected males tended to breed closer to the roost location. Male cockatoos are largely philopatric to their natal roost, and exhibit long-lasting foraging associations, preferentially with kin. Our results demonstrate carry-over between these social networks and the fine-scale arrangement of breeding individuals, consistent with an active choice to retain stable, across-context social relationships.

## Introduction

Social structures such as dominance hierarchies and social networks have been documented across diverse taxa and play an important role in regulating access to resources [1], facilitating information sharing [2], fostering cooperation [3], influencing health [4] and determining reproductive success [5]. In some species, these structures can be highly stable. However, recent evidence suggests that even in species that exhibit seasonal sociality, social associations can have carryover effects across seasons or contexts. For instance, zebra finches (*Taeniopygia guttata*) that breed synchronously exhibit stronger social ties in the subsequent non-breeding season [6], while territorial breeding great tits (*Parus major*) breed closer to those they previously associated with during the winter flocking period [7].

Retaining social associations across seasons can benefit individuals in various ways. For example, great tits that breed nearer to birds familiar from the previous winter or breeding-season [7, 8] continue to experience the benefits of that sociality, including access to social information about nesting material [9] and enhanced joint nest defence [10]. Additionally, if competition for sites limits access to preferred breeding locations [11], carry-over of dominance relationships should be more beneficial than re-establishing dominance hierarchies [12]. However, despite these benefits, social carry-over between seasons is not always observed. In migratory white-crowned sparrows, for example, winter social communities do not transition to breeding locations [13] despite stability between successive winters [14]. One potential explanation for these different outcomes might be that social carry-over is largely driven by co-occurrence in space [15, 16]. That is, if the spatial distribution of individuals in winter influences the arrangement of breeding territories, for example through spatial constraints, similar habitat preferences or patterns of space-use, then social associations may exhibit continuity between seasons even without any active preferences for prior social associates.

Study in species with communally roosting or colonially breeding social systems provides an opportunity to disentangle the influence of spatial and social decisions on social carry-over between seasons. In both cases, individuals share a single spatial location, either during breeding (in case of colonial breeders [17]), or during non-breeding periods (in communally roosting species [18]). This means that individuals effectively *‘reset’* by aggregating into one location, removing spatial constraints on the distribution of individuals from one season to the next. However to date, the potential for social carry-over effects in these species has been relatively less studied. Here, we investigate the relationship between social structures and breeding locations in one such species, a non-territorial, communally roosting parrot, the sulphur-crested (SC-) cockatoo (*Cacatua galerita*).

SC-cockatoos form year-round communal night roosts with fission-fusion foraging dynamics around the roost-site [19]. During spring, breeding pairs will split from this roost to occupy and defend nearby tree hollows, but remain non-territorial. Breeding pairs and their dependent offspring then return to the communal roost soon after fledging. Like most parrot species, SC-cockatoos are secondary cavity nesters that rely on natural hollow formation in large old-growth trees [20]. Suitable breeding hollows are therefore a limited resource, particularly in human modified environments [20]. This resource limitation creates a competitive environment that may shape both breeding behaviour and broader social dynamics.

Previous research on foraging associations in SC-cockatoos has found that individuals maintain stable social associations and linear dominance hierarchies that persist between successive nonbreeding seasons [21, 22]. However, it is unknown whether these social structures have consequences for breeding decisions. Our study therefore aimed to address two related questions about social carry-over. First, we examined whether there was a correlation between strength of dyadic associations in winter and spring and the spatial distribution of occupied nest hollows, hypothesising a positive correlation between breeding proximity and social association. Second, we investigated the relationship between breeding status, hollow occupation and dominance, hypothesising that individuals observed breeding will be higher ranked in the dominance hierarchy, and that higher-ranked individuals will occupy hollows closer to the communal roost site.

## Methods

### (a) Study site and subjects

The study was conducted in two neighbouring roost-sites in north Sydney, Australia: Balmoral Beach (BA) and Clifton Gardens (CG; Table S1). At each site, individuals were habituated to the observer, before being individually marked using non-toxic dye (Marabu Fashion Dye, MARABU GmbH) using methods detailed in Penndorf et al. 2023 [19]. As part of a parallel project, birds were also habituated and marked at 3 roosts outside the focal study area [19]. In addition to the paint-marked individuals, 144 birds had previously been wing-tagged as part of the citizen-science project Big City Birds [21, 23]. Age (juveniles: <7 years, adults: >7 years) and sex (only for adults) were assessed by eye-colour.

### (b) Data collection

Social data was collected close to the roost location (300-680m distance) over two 10-day periods in winter (July 8th-20th) and spring (September 19th-October 2nd 2019). During each daily session (July: 3h, September: 2.5h), presence scans [24] were conducted every 10 minutes to record the identities of all individuals present. Between scans, aggressive interactions were recorded using all occurrence sampling [24]. In addition, opportunistic scans were conducted at one intermediate foraging site, where birds of the two focal roosts mixed on a regular basis when fed by locals. Altogether, the final dataset included 511 scans of 375 individuals (July: 227 scans of 313 individuals, September: 284 scans of 239 individuals).

Between August and October 2019, random transects were conducted in a 1 km radius around each of the roost sites. As SC-cockatoos tend to display in front of their hollow around sunrise and sunset prior to initiating breeding, transects were conducted during these time periods, but opportunistic observations during the day were also included. We identified hollows either by sound (displays) or visually, and then recorded (i) the location of the hollow, and (ii) the identity of the bird(s) observed entering. To differentiate between prospecting individuals and breeders, identities were only considered as confirmed if a bird had been observed entering or emerging from a hollow on at least three separate occasions.

### (c) Social metrics

To create association networks, we created two group-by-individual matrices (one for the prebreeding period, one for the breeding period). Individuals were assigned to the same group if they had been observed in the same presence scans. As estimates for rarely observed individuals may be inaccurate [25], we thresholded the data to include only individuals present in at least six scans (detailed in [19, 26]). We then used a simple ratio index (SRI [27], R-package *asnipe* [28]) to create two undirected association networks (pre-breeding season and breeding season). The final association networks included 196 (July) and 155 (September) individuals.

For each roost location, we calculated a separate dominance hierarchy for July (pre-breeding season) and September (breeding season) using randomised Elo-rating (R-package *aniDom* [29], sigma=1/300, K=200, randomisations=10,000), including only individuals with seven or more aggressive interactions during each period at the specific roost location [22]. This resulted in 94 (July) and 75 (September) individuals being included in the dominance hierarchy in BA, and 67 (July) and 78 (September) individuals in the dominance hierarchy in CG.

### (d) Data analysis

To test whether social associations were correlated with hollow proximity, we calculated a proximity index as follows:

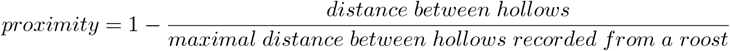

Therefore, our proximity measure could range between 0 (maximal distance between hollows recorded from one roosting group) and 1 (hollows in the same tree).

We compared hollow proximity with social association strength in the winter and spring social networks using multiple regression quadratic assignment procedures (MRQAP) via the *asnipe* package [25], where the significance was ascertained using permutations [25] and roost comembership was included as a covariate. Separate models were run for males and females.

To test the putative effect of rank or social network centrality on whether a bird was observed breeding, we ran a binomial generalized linear mixed model for each sex and social data observation period, examining whether a bird was observed at a hollow (binary, 0/1), as function of the standardized within-sex and roost dominance rank (continuous, 0-1), summed association strength, and the number of scans in which an individual was recorded (to control for potential variation in observability or residency), and roost (categorical, BA or CG).

Finally, to test whether higher-ranked or more socially central individuals tended to occupy hollows closer to the roost site, we ran a linear mixed model with distance in meters from roost as the response variable and the same predictor variables of standardized within-sex and within-roost dominance rank, summed association strength, the number of scans in which an individual was recorded, and roost (categorical, BA or CG). As above, models were run separately for males and females, and for the spring and winter social datasets.

## Results

We located 45 breeding hollows (figure 1a) across the two study locations (Figure 1b, 1c). For 29 of these hollows, we could identify at least one of the breeders (*N*_*pair*_=12, *N*_*female only*_ = 7, *N*_*male only*_= 10, Table S1). This resulted in 29% of adults observed breeding at BA, and 44% at CG, which is in line with previous estimates in urban populations of SC-cockatoos of approximately 30% of adults actively breeding [20].

**Figure 1.**
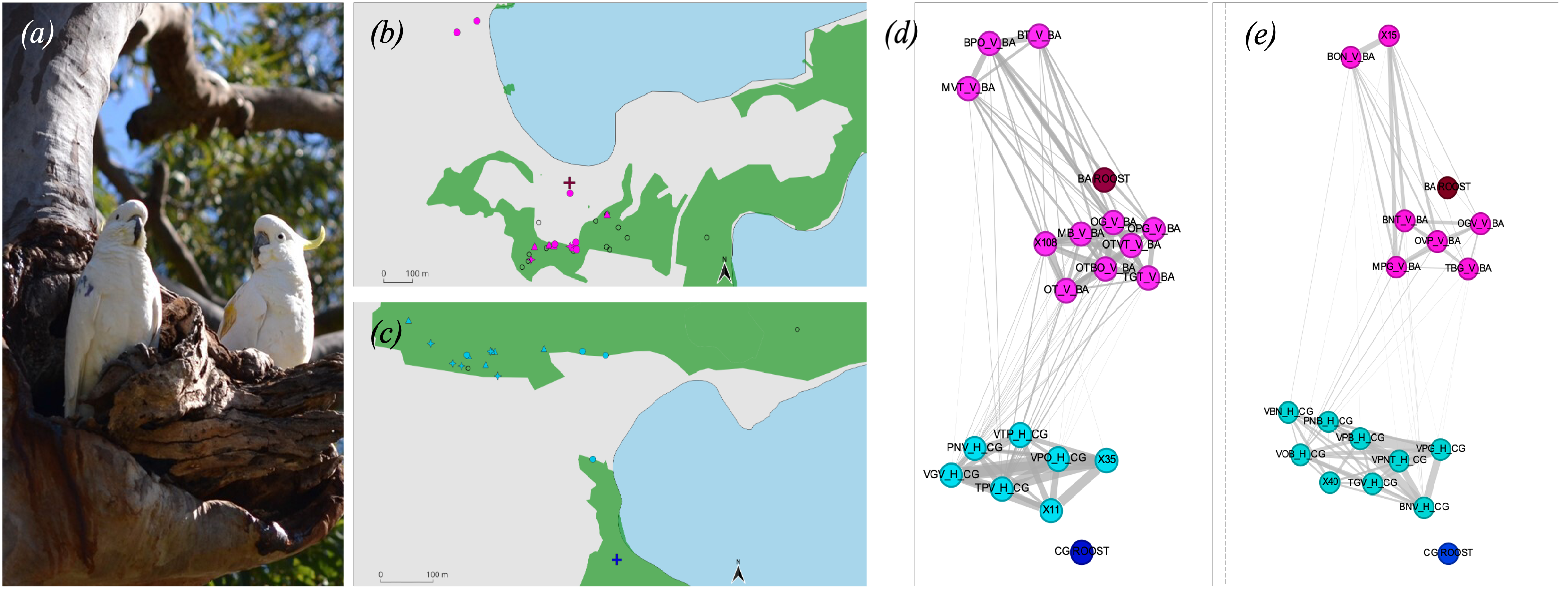
Location of hollows and social network ties between individuals holding hollows. (a) Photograph showing a breeding pair of SC-cockatoos—X11 (male) and BNV (female)—at their nest hollow. (b) Map of Balmoral Beach roost site, and (c) map of Clifton Gardens roost site. Cross marks roost site, coloured circles are hollows where the ID of both breeding individual was known, triangles are hollows where the ID of the breeding male was known, diamonds are hollows where the ID of the breeding female was known, and empty circles are hollows where neither individual was identified. Breeding males (d), but not females (e) exhibit stronger social associations with breeding neighbours. Social networks are summed from winter and spring data collection periods, and show roost members of Balmoral Beach in pink (roost site in dark red) and Clifton Gardens in cyan (roost site in dark blue). The relative position of nodes represents the geographic location of occupied hollows, and the edge weight represents association strength between individuals.

**Figure 2.**
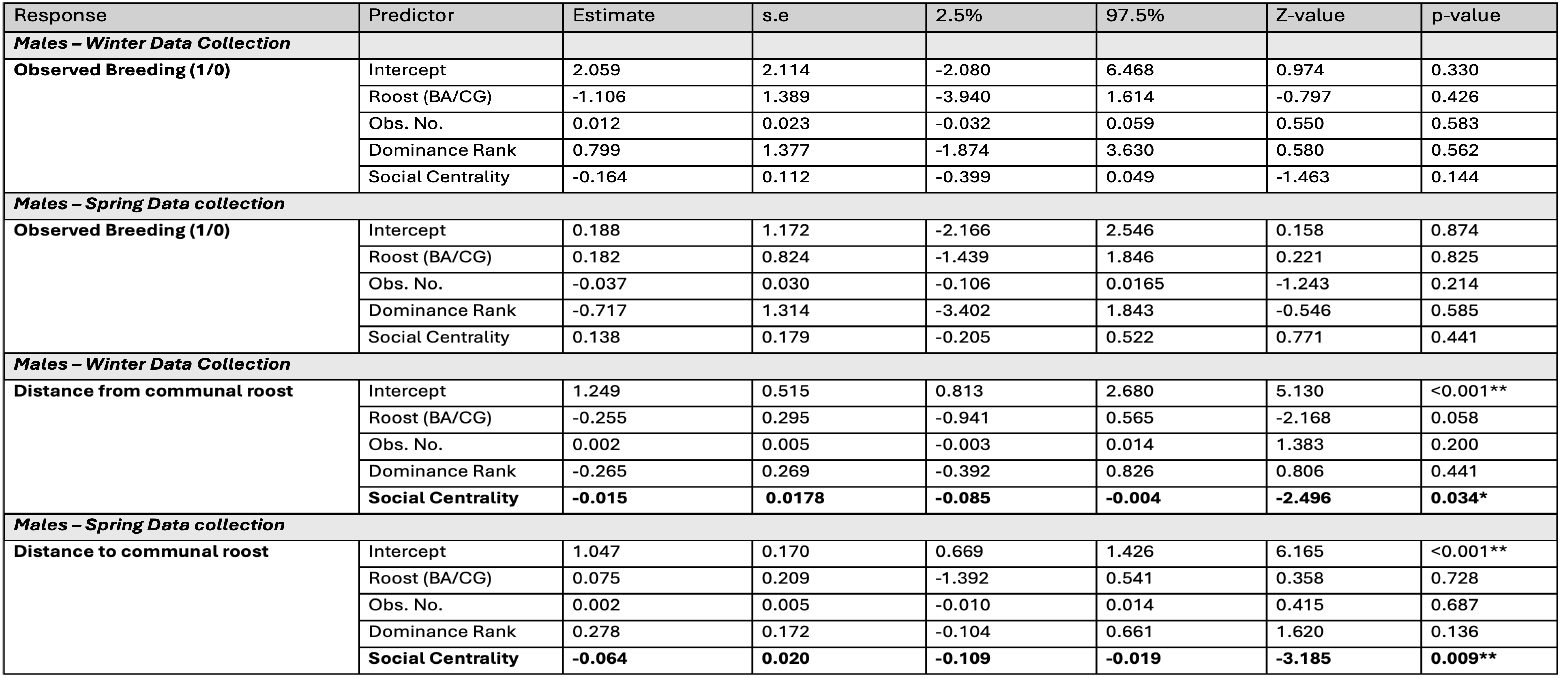
Statistical test results for males of general linear model of observed hollow occupation (1/0) and linear mixed model of hollow distance to the communal roost site with independent variables of with Dominance Rank, Social centrality (association strength) and extent of observation during social scans (Obs. No.). Significant p-values (< 0.05) are given in bold. Results for females are given in supplementary material.

As expected from previous work showing social stability in SC-cockatoos, there was a positive correlation between the winter and spring social networks (mantel test: r=0.65, p<0.001), and between the winter and spring dominance hierarchies.

Hollow proximity was correlated with social association strength in males, but not in females, whereby males that bred closer together associated more both in the winter social network (males: n=18, es=0.71, p=0.005; females: n=15, es=0.29, p=0.436) and in the spring social network (males: n=14, es=1.16, p=0.002; females: n=13, es=0.13, p=0.75): Figure 1d, Table S1.

There was no effect of social network centrality on the likelihood of being observed breeding in males (winter: es=-0.16, p=0.14; spring: es=0.14, p=0.44). However, more socially connected males were more likely to occupy hollows closer to the communal roost site, with this effect observed in both the winter and spring social networks (winter: es=-2.50, p=0.03; spring: es=-3.18, p=0.009). This effect was not driven by observability, as the number of social scans in which an individual was recorded was not related to the distance of their hollow to the roost (winter: es=0.002, p=0.20; spring: es=0.002, p=0.69): Table 1. Social network centrality in females was unrelated to the likelihood of breeding (winter: es=-0.05, p=0.72; spring: es=1.14, p=0.52) or distance to roost (winter: es=-0.04, p=0.58; spring: es=-0.01, p=0.83): Table S2.

In males, there was no correlation between dominance rank and the likelihood of being observed breeding or distance of an individual’s hollow to the communal roost (winter: es=0.80, p=0.56; spring: es=-0.71, p=0.58; controlling for roost membership and observation number): Table 1. In females, there was no relationship between dominance rank and distance of the occupied hollow to the roost (winter: es=0.16, p=0.41; spring: es=-0.22, p=0.35). However there was a significant relationship between the dominance rank of females in spring and hollow occupancy (winter: es=-1.36, p=0.46; spring: es=-3.85, p=0.02): Table S2.

## Discussion

Our study uncovered a sex-dependent correlation between breeding decisions and social structure in wild SC-cockatoos. Males—but not females—associated more with neighbouring breeders of the same sex, and males that were more central in the social network tended to occupy hollows closer to the communal roost site. However, and contrary to our prediction, there was no relationship between higher dominance rank in males and either the likelihood of being observed breeding or of occupying breeding hollows close the roost. Altogether, these results suggest that there is a significant carry-over between foraging social networks amongst male SC-cockatoos and their choice of breeding locations, highlighting the importance of social bonds across different contexts.

We found that males breeding in proximity of each other were more likely to have been stronger associates during the previous winter, with these foraging associations then retained through the spring breeding period. Similar correlations have been described in other species. For example, in great tits, winter associations at feeders positively predicted nest proximity during the next breeding season [7]. This effect was seemingly long lasting, as, in turn, breeding position was correlated with the subsequent winter’s social associations and breeding proximity in the following spring [7]. However these studies have largely been conducted on species that form breeding territories, making it difficult to fully disentangle active social preferences from alternative mechanisms. For example, a preference to remain close to the territory in winter could lead to spatial assortment in the formation of winter flocks, or individuals’ preference to establish a territory on the foraging home-range shared by a winter flock could lead to them forming neighbouring breeding territories [7, 30]. By contrast, SC-cockatoos are non-territorial when breeding, with breeding locations only limited by the distribution of available hollows. At all other times of year, the entire local population shares a communal sleeping site. It therefore seems unlikely our observed correlation between male foraging associations and breeding proximity could arise through mechanisms other than active social choice, especially given that this effect was only observed in one sex.

It is interesting to consider why we observed sex differences in social carry-over effects. One potential explanation could be sex-specific variation in the acquisition of the breeding hollow or in breeding behaviour. For example, juvenile male song sparrows *Melospiza melodia* tend to establish their breeding territories near to males from first-year foraging flocks [30]. Yet while more research is needed, pairs of SC-cockatoos appear to mostly gain breeding opportunities through aggressive take-over of hollows from other pairs, with observations suggesting that these contests involve both sexes (*pers. obs*.). Both sexes also display at the hollow, and incubate and feed chicks. However one important difference might be that, at night, females sleep in the hollow while males sleep outside in the same or nearby trees. Familiarity with neighbours might therefore allow males to retain some of the benefits of communal roosting at a smaller scale while breeding.

Another potential explanation could be to do with sex differences in sociality more broadly. Male SC-cockatoos tend to be more philopatric than females, with many remaining in their natal roost [19]. While we did not measure kinship, a previous study in this population has also shown that individuals preferentially socially associate with kin [19]. It therefore seems likely that nearby breeding males may also be more related, increasing the potential benefits of neighbour familiarity even further.

It is important to note that given the between-season social stability in this system, we cannot disentangle the directionality of this relationship; males may form stronger associations with individuals they breeding near to, or prefer to breed nearer to those with whom they share stronger social bonds. However, irrespective of directionality, familiarity with breeding neighbours could result in benefits. For example, great tits that share territorial boundaries with familiar individuals experience increased breeding success [31, 10, 32]. The potential benefits of social carry over in this system remain to be explored, but could include shared defence of the hollow; for example against nest predators (e.g. brushtail possums *Trichosurus vulpecula* and monitor lizards *Varanus spp*.), or from hollow take-over by conspecifics. Alternatively, it could include the roosting benefits as discussed above, including shared vigilance for nocturnal predators (Powerful owls *Ninox strenua*).

Interestingly, despite not all adult birds being observed breeding, and observations of contests at hollows (*pers. obs*.), there was no relationship between dominance rank in males and either the likelihood of being observed breeding or of occupying breeding hollows close the roost. Yet there was an effect of social centrality, with central individuals tending to breed closer to the communal roost site. We consider it unlikely that this result is simply due to increased observability, as 1) it was only observed in males, 2) it was consistent across both winter and spring social networks, and 3) hollow distance was not correlated to the total number of observations. Yet similarly to above, we cannot distinguish directionality: more social individuals may choose to breed closer to the communal roost site, or, if breeding hollows close to the roost site are preferred, more socially central individuals may have an advantage in gaining these hollows. One intriguing possibility that arises from our results is that social connections may be more important in successfully gaining a hollow than dominance rank, at least in males. However, we cannot exclude the possibility that success at gaining a breeding hollow is more related to female dominance, to a pair-level metric of dominance, or an aspect of competitive ability not measured in our study.

Finally, in contrast to the results in males, breeding females tended to more high ranking within the dominance hierarchy of their own sex. This result is consistent with findings in other bird species, where dominant females are more likely to breed [33]. However, we can only speculate on whether only high ranking females were able to obtain hollows, or whether breeding status subsequently affected the dominance status of females. Previous studies on spectacled parrotlets (*Forpus conspicillatus*) demonstrated that breeding pairs became dominant over non-breeding pairs only after occupying a cavity [34], an effect consistent with our observation that the link between female dominance and breeding status was only observed in spring. However, in this case, we would also expect breeding males in our study to occupy high dominance ranks. One potential explanation is that breeding status has a more pronounced effect on the female dominance rank compared to males. A second explanation is that more dominant females are successful in “breaking up” existing pairs by displacing the other—lower ranking—female to take her place, a behaviour previously described in spectacled parrotlets [34]. A third explanation is that males may prefer high-ranking females. Mate preferences for high-ranking individuals have been shown in several species, although often in the opposite direction. For example, in black-billed magpies (*Pica pica*) and spectacled parrotlets females—-but not males—preferentially formed pairs with high-ranking mates [34]. Further studies will be needed to elucidate the drivers of this relationship.

In conclusion, we found a sex-dependent relationship between breeding proximity and social association in wild SC-cockatoos. Males—but not females—tended to breed in closer proximity to individuals with whom they shared stronger foraging associations. As foraging associations were measured at the communal roost site, and individuals are non-territorial, this suggests an active social choice to carry social associations over between these two contexts, with this interpretation strengthened by our observation of this result only in males (the more philopatric sex [35]). Although we could not disentangle directionality in our study, it adds to the growing picture of strong social stability in this long-lived species [21, 22], despite large group sizes and daily fission-fusion social dynamics [19]. Overall, our study contributes to disentangling the intricate relationship between spatial structure and social networks in wild populations.

## Ethics statement

Research was approved by the Ethics Council of the Max Planck Society, Germany (application no. 2018_12). All procedures were approved by the ACEC (ACEC Project No. 19/2107), and were conducted under a NSW Scientific License (SL100107).

## Authors’ contribution statement

LMA secured funding. JP and LMA conceived of the study, curated the data and conducted statistical analysis and visualisation. JP undertook fieldwork. JP and AC wrote the paper. All authors discussed the results and edited the manuscript.

## Acknowledgements

We would like to acknowledge the Gamaragal and Gadigal people as the traditional custodians of the land on which this study was conducted.

We thank John Martin for his help and guidance in the field, and to the rest of the Aplin lab group for helpful discussions.

