## Appendix for "Sex-dependent carry-over between winter associations and breeding proximity in a communally roosting parrot"

#### Electronic Supplementary Materials

**Manuscript title:** Sex-dependent carry-over between winter social associations and breeding proximity in a communally roosting wild parrot

**Authors:** Julia Penndorf, Annegret Csak, Lucy M. Aplin

#### Tables

- **Table S1**: Location of roosting locations.

#### Figures

- **Figure S1**: Dominance ranks of each member of identified breeding pairs at the two roosts included in the study.

### S1 Supplementary Tables

Table S1: **Location of the two roosting locations considered in the study, their estimated size (number of individuals) and the number of nearby hollows.** mf: the identity of both individuals are known, m: only the identity of the breeding male is known, f: only the identity of the breeding female is known, u: the identities of both individuals could not be assessed with certainty

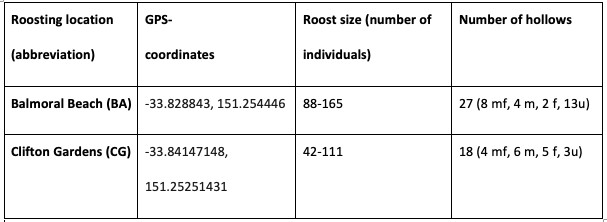

### S2 Supplementary Figures

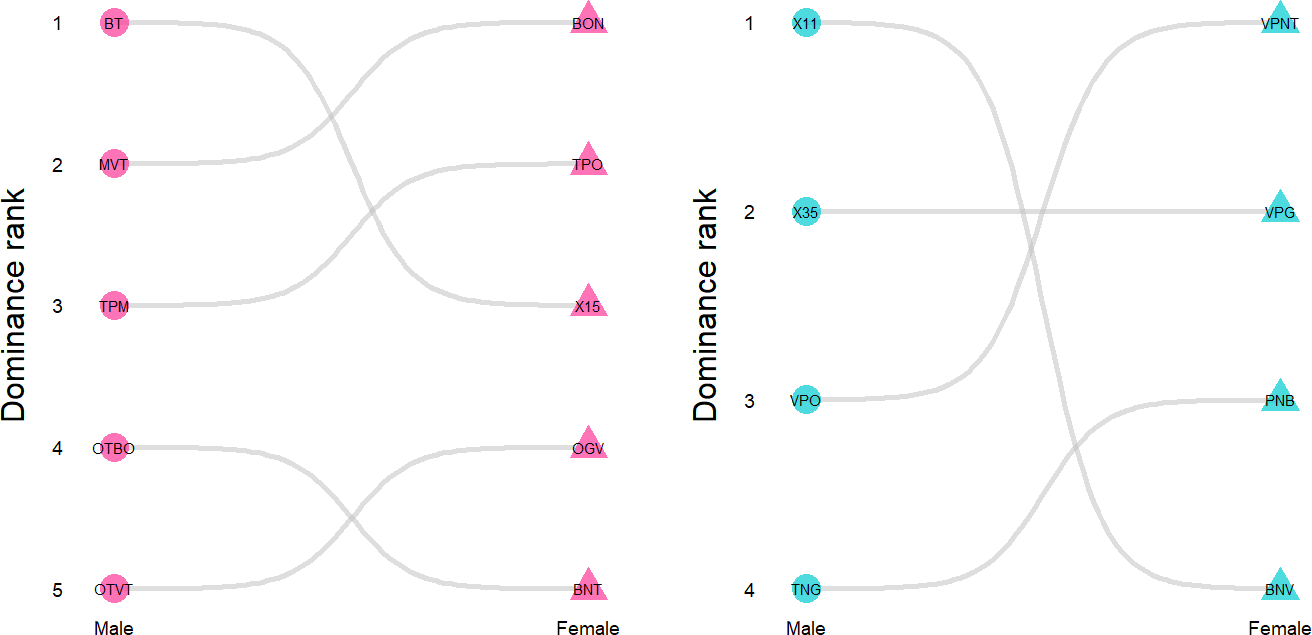

Figure S1: Dominance ranks of each member of identified breeding pairs at the two roosts included in the study: BA (pink, left graph) and CG (blue, right graph). Hierarchies in SC-cockatoos are sex-segregated; therefore we ranked individuals within sex. Males are represented by a circle, females by a triangle. The grey edges link members of the same breeding pair.
